# Norepinephrine and glucocorticoids modulate chronic unpredictable stress-induced increase in the type 2 CRF and glucocorticoid receptors in brain structures related to the HPA axis activation

**DOI:** 10.1101/2021.02.02.429424

**Authors:** Marilia B Malta, Joelcimar Martins, Leonardo S Novaes, Nilton B dos Santos, Luciane Sita, Rosana Camarini, Cristoforo Scavone, Jackson Bittencourt, Carolina D. Munhoz

## Abstract

The stress response is multifactorial and enrolls circuitries to build a coordinated reaction, leading to behavioral, endocrine, and autonomic changes. These changes are mainly related to the hypothalamus-pituitary-adrenal (HPA) axis activation and the organism’s integrity. However, when self-regulation is ineffective, stress becomes harmful and predisposes the organism to pathologies. The chronic unpredictable stress (CUS) is a widely used experimental model since it induces physiological and behavioral changes and better mimics the stressors variability encountered in daily life. Corticotropin-releasing factor (CRF) and glucocorticoid (GCs) are deeply implicated in the CUS-induced physiological and behavioral changes. Nonetheless, the CUS modulation of CRF receptors and GR and the norepinephrine role in extra-hypothalamic brain areas were not well explored. Here, we show that 14-days of CUS induced a long-lasting HPA axis hyperactivity evidenced by plasmatic corticosterone increase and adrenal gland hypertrophy, which was dependent on both GCs and NE release induced by each stress session. CUS also increased CRF_2_ mRNA expression and GR protein levels in fundamental brain structures related to HPA regulation and behavior, such as the lateral septal nucleus intermedia part (LSI), ventromedial hypothalamic nucleus (VMH), and central nucleus of the amygdala (CeA). We also showed that NE participates in the CUS-induced increase in CRF_2_ and GR levels in the LSI, reinforcing the *locus coeruleus* (LC) involvement in the HPA axis modulation. Despite the CUS-induced molecular changes in essential areas related to anxiety-like behavior, this phenotype was not observed in CUS animals 24 h after the last stress session.

**Highlights:**

- CUS persistently increased plasma CORT levels via GCs and NE signaling.
- CUS persistently increased CRF_2_ mRNA in extra-hypothalamic brain areas.
- CUS increased GR protein levels in brain regions related to GCs release control.
- NE and GCs participate in the CUS-induced increase in CRF_2_ and GR levels.
- LSI could be the brain nucleus that dictates the fine-tuned response of CUS.
- CUS animals did not present anxiety-like behavior.

## 1. Introduction

Physical and psychological stressors recruit different brain nuclei to build a coordinated stress response that activates the neuroendocrine and autonomic systems and causes behavioral changes [1]. Such response enfolds a conserved and efficient interconnection of several systems to maintain the organism’s integrity [2]. However, when this self-regulation is ineffective, stress becomes harmful and predisposes the body to diseases, including cardiovascular, immune, and psychiatric disorders [3,4].

Several brain regions, such as the paraventricular nucleus of the hypothalamus (PVH), septal-hippocampus complex, amygdalar nuclei, pre-frontal and cingulate cortices, *locus coeruleus* (LC), and parabrachial and raphe nuclei orchestrate the stress response [5]. The converging signals activate the hypothalamus-pituitary-adrenal (HPA) axis. The PVH releases the corticotropin-releasing factor (CRF), which results in the synthesis and release of the adrenocorticotropic hormone (ACTH) by the anterior pituitary, ultimately leading to glucocorticoids (GCs) synthesis and release by the adrenal glands [6–8].

The CRF actions in the central nervous system (CNS) and the periphery are critical to integrating and coordinating several physiological systems. During stress adaptation, these complex responses are fine-tuned by several CRF related peptides, including the urocortins (UCNs), that exert complementary or sometimes contrasting actions to CRF itself [9]. The CRF peptides (CRF and urocortins) act by binding to two different G-protein coupled receptors, the CRF type 1 and type 2 receptors (CRF_1_ and CRF_2_), which play different roles in stress responses. The outcomes of CRF receptors activation depend on the ligands and brain nuclei and decisively dictate the stress-induced behavioral state [reviewed on [10–13]]. For example, CRF_1_ activation has been associated with the endocrine effects of CRF and behavioral response to stress, whereas the role of CRF_2_ is essential to enable physiological and psychological homeostasis and oppose the initial stress response linked to CRF_1_ activation [14,15,10–12,16,17,13]. Particularly, the CRF_2_ receptor’s genetic deficiency increases anxiety-like behavior and the sensitivity to stress [14] but attenuates the responses to the stress induced by opiate withdrawal [18]. Thus, the contribution of CRF_2_ on responses to different stressful events warrants investigation.

In addition to the HPA axis activation, CRF also modulates the stress-induced sympathetic response through the recruitment of the LC, adrenal medulla, and peripheral sympathetic system and plays a significant role in integrating the central and peripheral norepinephrine (NE) release secondary to stressful events [reviewed on [12]], which is fundamental to mediate the neuroendocrine changes and to bring together the stress perception and physiological and behavioral alterations.

However, acute and chronic stress can use different pathways to change the control of the HPA axis and autonomic responses [19,1,5]. In a chronic stress situation, neurochemical evidence suggests a continued increase in the HPA and adreno-medullary axis excitability in addition to an augmented sensibility of the LC to CRF and to the enzyme tyrosine hydroxylase, which elevates NE synthesis (CRF-NE loop) [9]. Some of these changes could be associated with the loss of GCs self-regulation in the pituitary and PVH. Furthermore, it was also reported neuronal atrophy in the hippocampus and frontal cortex, as well as an increase in the CRF expression in the central nucleus of the amygdala (CeA) and autonomic nuclei, which resulted in the HPA axis hyperactivity and increased excitability of PVH [20].

The disruption of the GCs negative feedback on the HPA axis results in its sustained activation, causing the maintenance of elevated systemic GCs levels. Consequently, the higher availability of these hormones allows them to reach brain structures such as the amygdala and the LC, whose projections reinforce the HPA axis activation and promote changes in behavior and normal physiology [19,21,22]. In fact, due to their pleiotropic action, the sustained increase in the systemic levels of GCs can trigger metabolic diseases, immunosuppression and contribute to the development of autoimmune diseases and mood disorders [23].

The chronic unpredictable stress (CUS) paradigm is a well-accepted experimental model of stress-induced brain plasticity and mood disorders, such as depressive-like behaviors [24]. The mixture of psychological and physical stressors, along with reduced chances of adaptation to the different stress stimuli, present significant similarity to stressors encountered in daily life [25,26]. Considering previous data of loss of HPA axis self-regulation in animals submitted to 14 days of CUS [27] and the few evidence of the NE role in CUS studies, the present work evaluates the changes in the CRF_2_ and glucocorticoid receptors (GR) in the central nervous system (CNS) of rats exposed to CUS, the influence of GCs and NE signaling in those molecular changes and their implications in CUS-induced anxiety-like behavior. We found that CUS increased CRF_2_ mRNA expression and GR levels in central brain structures related to the HPA axis, and both GCs and NE signaling modulated these effects. Interestingly, CUS animals did not show any anxiety-like behavior in EPM and OP tests.

## 2. Materials and Methods

### 2.1. Animals

Animals (Adult male Wistar rats, 60-days old at the beginning of the experiments) from the Facility for SPF Rat Production at the Institute of Biomedical Sciences - Animal Facility Network at the University of São Paulo were used following the standards of the Ethics Committee for Animal Use of the Institute of Biomedical Sciences/University of São Paulo (CEUA- ICB 102/06 and 75/05) and the guidelines of the Brazilian National Council for the Control of Animal Experimentation (CONCEA). All efforts were taken to minimize animal suffering and reduce the number of animals used to the minimum required for detection of significant statistical effects.

### 2.2. Chronic unpredictable stress paradigm

Adult male Wistar rats housed in groups of 3 or 4 with *ad libitum* rat chow and water were kept under a 12 h light/dark cycle. The animals were randomly assigned to either control (non-stressed) or chronic unpredictable stress (CUS) group and submitted or not to the different pharmacological treatments. The CUS protocol was based on [27]. Briefly, the animals were subjected to diverse psychological and physical stressors for 14 days, applied as follows: day 1 (2:00 p.m.) restraint, 60 min; day 2 (9:00 a.m.) forced swim, 15 min; day 3 (3:00 p.m.) cold isolation, 90 min; day 4 (7:00 p.m.) lights on, overnight; day 5 (10:00 a.m.) forced swim, 5 min; day 6 (7:00 p.m.) water and food deprivation, overnight; day 7 (2:00 p.m.) restraint, 120 min; day 8 (3:00 p.m.) light off, 120 min; day 9 (9:00 a.m.) forced swim, 5 min; day 10 (7:00 p.m.) lights on, overnight; day 11 (2:00 p.m.) cold isolation, 90 min; day 12 (9:00 a.m.) restraint, 60 min; day 13 (7:00 p.m.) water and food deprivation, overnight; day 14 (9:00 a.m.) restraint, 60 min. In all CUS experiments, stress stimuli were applied in the same order described above. All animals were studied 24 hours after the last stressor. The control rats were manipulated every day for 10 min in their home cages to avoid nonspecific handling effects.

### 2.3. Pharmacological treatments

Metyrapone (2-Methyl-1,2-di-3-pyridyl-1-propane; Sigma), a steroid 11β-inhibitor, was first dissolved in 100% ethanol and then diluted (1:10) in (NaCl 0.9%). Metyrapone (Met; 50 mg/kg) was administered intraperitoneally (i.p.) 30 min before each stress session according to [28]. We attempt to avoid a dose that could interfere with the circadian cycle of glucocorticoid synthesis.

Alpha and beta noradrenergic antagonists, phentolamine (P) and atenolol (A), respectively, were individually dissolved in 0.9% saline and concomitantly (AP) injected i.p. 20 min before each stress section in the doses of 5 mg/kg and 2.5 mg/kg respectively [29]. The treatment with both antagonists was chosen considering the presence, distribution, and action of alpha and beta noradrenergic receptors in the body and brain areas, which allowed us to blunt all noradrenergic influence regardless of whether central or peripheral.

For control of the respective pharmacological treatments, control animals were administered with Met or AP simultaneously as the CUS animals and returned to the animal facility.

### 2.4. Experimental groups and design

We randomly divided the animals into the following groups:

- Control: no stressed animals
- CUS: animals submitted to chronic unpredictable stress
- C + Met: control animals submitted to chronic metyrapone treatment
- C + AP: control animals submitted to chronic atenolol and phentolamine treatment
- CUS + Met: animals submitted to CUS and treated chronically with metyrapone
- CUS + AP: animals submitted to CUS and treated chronically with atenolol and phentolamine

### 2.5. Euthanasia and perfusion

Twenty-four hours after the last stress session, the animals were anesthetized with an excess of isoflurane and perfused via the ascending aorta with ≅100 mL of cold 0.9 % saline in 1 min, followed by ≅ 900 mL of 4 % formaldehyde in dH_2_O at 4 °C, pH 9.5, for approximately 25 min. After removed from the skull, the brains were post-fixed for 3 h in the same fixative solution plus 20 % sucrose and then transferred to RNAse free 20 % sucrose in phosphate buffer solution at 4 °C. Regularly spaced brain series (5 × 1-in-5) of 30 μm-thick frozen sections were cut in freezing microtome (SM2000R, Leica, Wetzlar, HE, Germany) in the coronal plane, collected in ethylene glycol-based cryoprotectant solution, and stored at −20 °C until assayed. To preserve RNA integrity for in situ hybridization all tubes and solutions used were RNAse Free all solutions were prepared fresh with DNAse and RNAse free reagent and miliQ water.

### 2.6. Plasma CORT concentration

Plasma corticosterone (CORT) was measured using an enzyme-linked immunoassay (Enzo Life Sciences). Before the saline perfusion, cardiac blood was collected, transferred to conical tubes (with heparin 10% v/v), and centrifuged at 3500 rpm for 15 min to obtain plasma samples. CORT titers were assessed following the manufacturer’s instructions. Plasma samples were diluted at 1:30 in assay buffer before use.

### 2.7. *In situ* hybridization for the CRF_2_ in rat brain

The *in-situ* hybridization for CRF_2_ was carried out using a ^35^S-labeled antisense cRNA probe from plasmid kindly provided by Paul Sawchenko (The Salk Institute for Biological Studies, La Jolla, CA, USA). Techniques for probe synthesis, hybridization, and autoradiographic localization of mRNA signal were adapted from [30]. In brief, brain sections were mounted onto electrostatically charged slides (Superfrost plus®, Fisher Scientific, USA) and digested with 10 μg/ml of proteinase K for 30 minutes at 37 °C. The labeled probes (sense and antisense) were used at concentrations of 10^6^ cpm/ml and applied to sections overnight at 56 °C in a solution containing 50% formamide, 0.6M NaCl, 10 mM Tris, pH 8.0; 1mM EDTA, 0.05% tRNA, 10 mM dithiothreitol, Denhardt’s solution, and 10% dextran sulfate. They were then treated with 20 μg/ml of ribonuclease A for 30 min at 37 °C and washed in 15 mM NaCl/1.5 mM sodium citrate at 55 – 60 °C. Sections were dehydrated and exposed to x-ray films for 1–2 days. The slides were then dipped in Kodak NTB-2 liquid autoradiographic emulsion, dried, and exposed at 4°C in the dark for 20 days. They were then developed with Kodak D-19 and counterstained with thionin (Nissl method) for anatomical reference purposes. Sections were dehydrated, defatted, and coverslipped with DPX mounting medium (Aldrich, Milwaukee, WI, USA). For CRF_2_ mRNA expression, the semiquantitative densitometric analysis was performed on emulsion-coated sections. Slides were coded and analyzed blinded. Densitometric analysis of the images was performed under dark-field illumination with the Image-Pro Plus software. We evaluated the CRF_2_ mRNA expression in the lateral septal nucleus intermediate part (LSI), and ventromedial hypothalamic nucleus (VMH) integrated optical density (IOD) in arbitrary units (a.u.). The results are expressed by mean of IOD ± SEM (a.u., n=6/group) at medial levels of LSI and VMH (0.20 and −3.14 mm from Bregma for LSI and VMH, respectively).

### 2.8. Immunohistochemistry for GR

Immunohistochemistry was performed using a conventional avidin-biotin immunoperoxidase protocol [31] and Vectastain Elite reagents. The free-floating sections were pretreated with hydrogen peroxide, blocked in 3% normal donkey serum, and incubated overnight at room temperature with anti-GR polyclonal primary antisera raised in rabbit (1:3000 sc-1004 GR Antibody (M-20) Santa Cruz). Sections were incubated for 1h in biotin-conjugated IgG donkey anti-rabbit (1:800, Jackson Laboratories) and 1h in avidin-biotin complex (1:333, kit elite Vector Labs). The peroxidase reaction was performed using 0.05% DAB and 0.025% nickel sulfate as chromogens, and 0.03% hydrogen peroxide. Sections were mounted onto gelatin-coated slides, dehydrated, defatted, and coverslipped with DPX mounting medium (Aldrich, Milwaukee, WI, USA). To evaluate the GR levels, we quantified the stained cells in the LSi, CeA, bed nucleus of the *stria terminalis* (BST), and PVH. The results are expressed as a mean of GR immunoreactive cells ± SEM. The total area was defined by the product between the number of sections analyzed and the area in a 20 X magnification for each nucleus (0.278 mm^2^ for LSi and CeA; 0.06 mm^2^ for and 0.04 mm^2^ for PVH).

For the CRF_2_-GR co-localization experiments, the immunohistochemistry for GR was performed as described and after the peroxidase reaction, the sections were mounted onto electrostatically charged slides (Fisher Superfrost Plus®, Fisher Scientific, USA). The *in-situ* hybridization for CRF_2_ was performed as described earlier.

### 2.9. Behavioral studies

To correlate the CUS-induced biochemical changes to the behavioral phenotype, the animals were examined for anxiety-like behavior using the elevated plus maze (EPM) and open field (OF) tests 24 hours after the last stress session in a non-aversive environment and after the acclimatization period.

The EPM apparatus consisted of two closed and two open arms with a central area (100 cm^2^) where the animals were positioned at the beginning of the test. The apparatus was made of dark blue painted wood elevated 65 cm from the floor and located in a soundproof test room under indirect light. A video camera was located above the center of the maze to record the rats. Anxiety-like behavior was assessed as a function of the decreased open arm exploration, measured by the total percentage of time at the open arms.

The open field (OF) apparatus consists of a circular arena (60 cm diameter) with opaque walls (50 cm high). The floor was divided into squares, which allowed the discrimination of two compartments, the central and peripheral. Each rat was placed on the periphery of the arena, and its activity was recorded for 5 min. This test was used to evaluate the anxiety-like behavior based on rodents’ preference to spend a significantly higher amount of time exploring the periphery of the arena than the unprotected center area.

In both tests, the animals explored the apparatuses for 5 minutes. The behavior was evaluated by the EthoVision System (Noldus Information Technology). Results are expressed as mean of percentage of the time that each group spent in the arms ± SEM (%) for the EPM and as a mean of the time in central zone ± SEM (%) for the OF. Different groups of animals were used for each test, and they were not previously exposed to neither EPM nor OF. All trials were videotaped, and the apparatus was cleaned with 5% (vol/vol) ethanol after each trial to avoid olfactory clues.

### 2.10. Statistical analysis

The statistical analysis results are expressed as mean ± SEM (n=6/group) and were calculated using the GraphPad Prism Software, version 5.0. Under the guidance of the statistical team of the Institute of Mathematics and Statistics (IME-USP), we adopted the Kruskal Wallis test [32] to analyze the results from *in situ* hybridization (CRF_2_ mRNA expression) and immunohistochemistry (GR levels). We used the Student T test to analyze the behavior. Values of p<0.05 were considered statistically significant.

## 3. Results

### 3.1. CUS persistently increased plasma CORT levels through GCs and NE signaling

CUS augmented plasma CORT levels 24 h after the last stress session (14.9 ± 2.2 μg/dL, p <0.05) when compared to control rats (4.9 ± 0.95 μg/dL) (Figure 1). This effect was accompanied by the hypertrophy of the adrenal gland (17.4 ± 0.8 and 14.2 ± 0.8% of adrenal gland CUS and control, respectively, p <0.05) in the CUS animals. Chronic treatment with Met (CUS + Met., 2.6 ± 0.7 μg/dL) and AP (CUS + AP, 3.9 ± 1.8 μg/dL) blunted the increase of CORT levels in CUS animals without altering CORT basal levels of control, non-stressed group in the same time point (4.9 ± 1.0 μg/dL, p<0.05). At 72 h-post-stress, both CORT levels and adrenal glands were no different from control.

**Figure 1.**
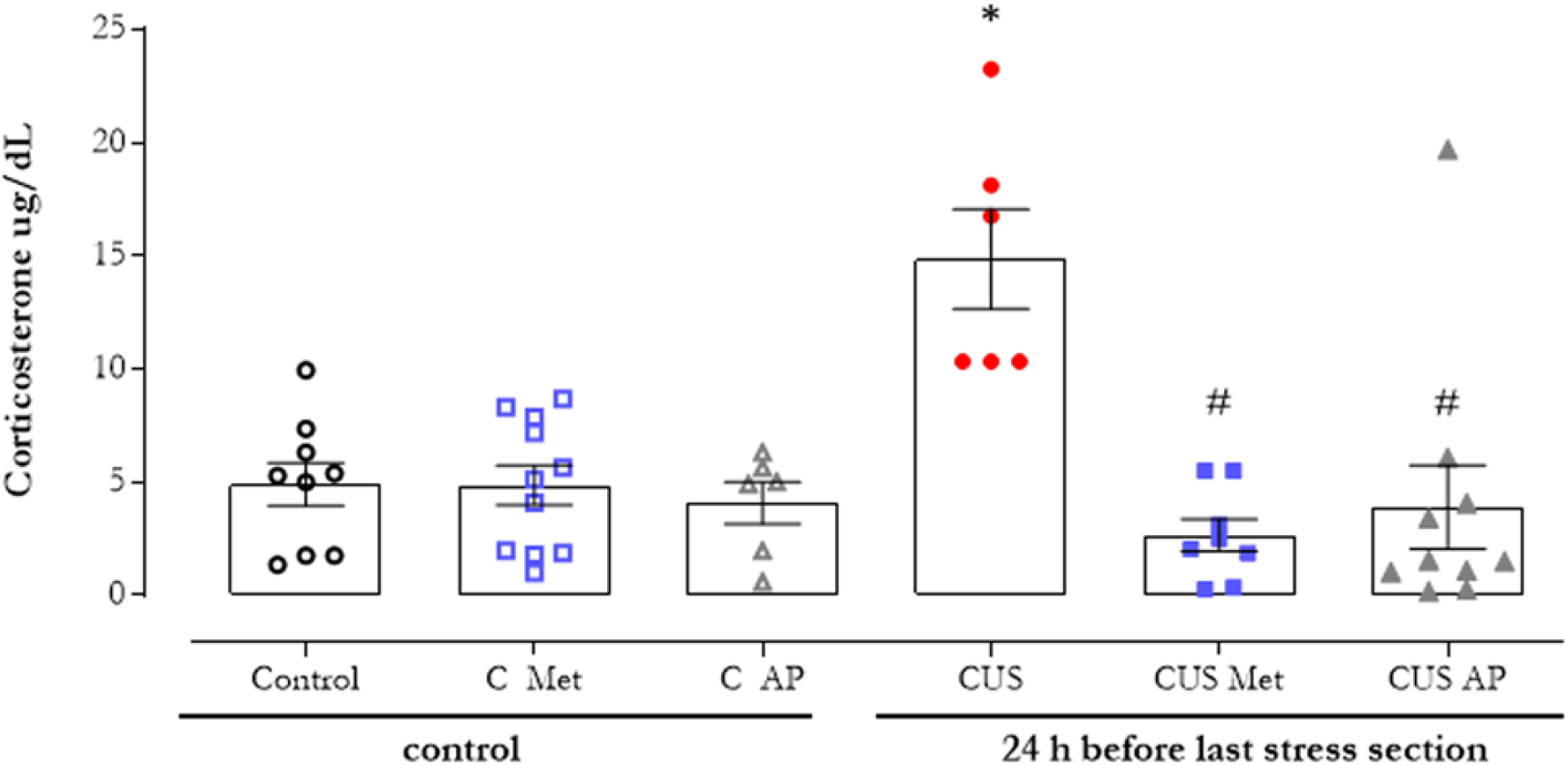
CUS-induced persistently increase in plasma CORT levels is modulated by GCs release and NE signaling. Plasma CORT from control and 14 days of chronic unpredictable stress (CUS) exposed animals treated or not with metyrapone (Met, 50 mg/kg, i.p., 30 min before each stress session) or atenolol and phentolamine (AP, 5 mg/kg and 2.5 mg/kg respectively, i.p., 20 min before each stress session) were collected 24-hours after the last stress session. Results are presented as mean ± SEM. Significance differences between groups are indicated as * p < 0.05 vs control; • p < 0.05 vs CUS, Kruskal Wallis and Mann Whitney tests (n = 6 for each group).

### 3.2. CUS persistently increased CRF_2_ mRNA expression in the LSi and VMH, an effect dependent on GCs and NE signaling

Using the *in-situ* hybridization assay, we observed an increase of the CRF_2_ mRNA expression in the LSI and VMH (1.4×10^6^ ± 1.3×10^5^ a.u. and 2.4×10^6^ ± 2.2×10^5^ a.u. for LSI and VMH, respectively) of CUS animals when compared to the control group (1.0×10^6^ ± 9.4×10^4^ a.u. and 7.9×10^5^ ± 1.0×10^5^ a.u. LSI and VMH respectively, p < 0.05) (Figure 2 and Table 1). Because GC levels were elevated 24 h after the last stress session, we tested whether this sustained CORT increase helped to mediate the effects of stress on CRF_2_ mRNA expression. As shown in figure 2, blunting the stress-induced GCs increase with metyrapone (Met) reduced the CRF_2_ mRNA expression in both LSI and VMH compared to CUS animals.

**Figure 2.**
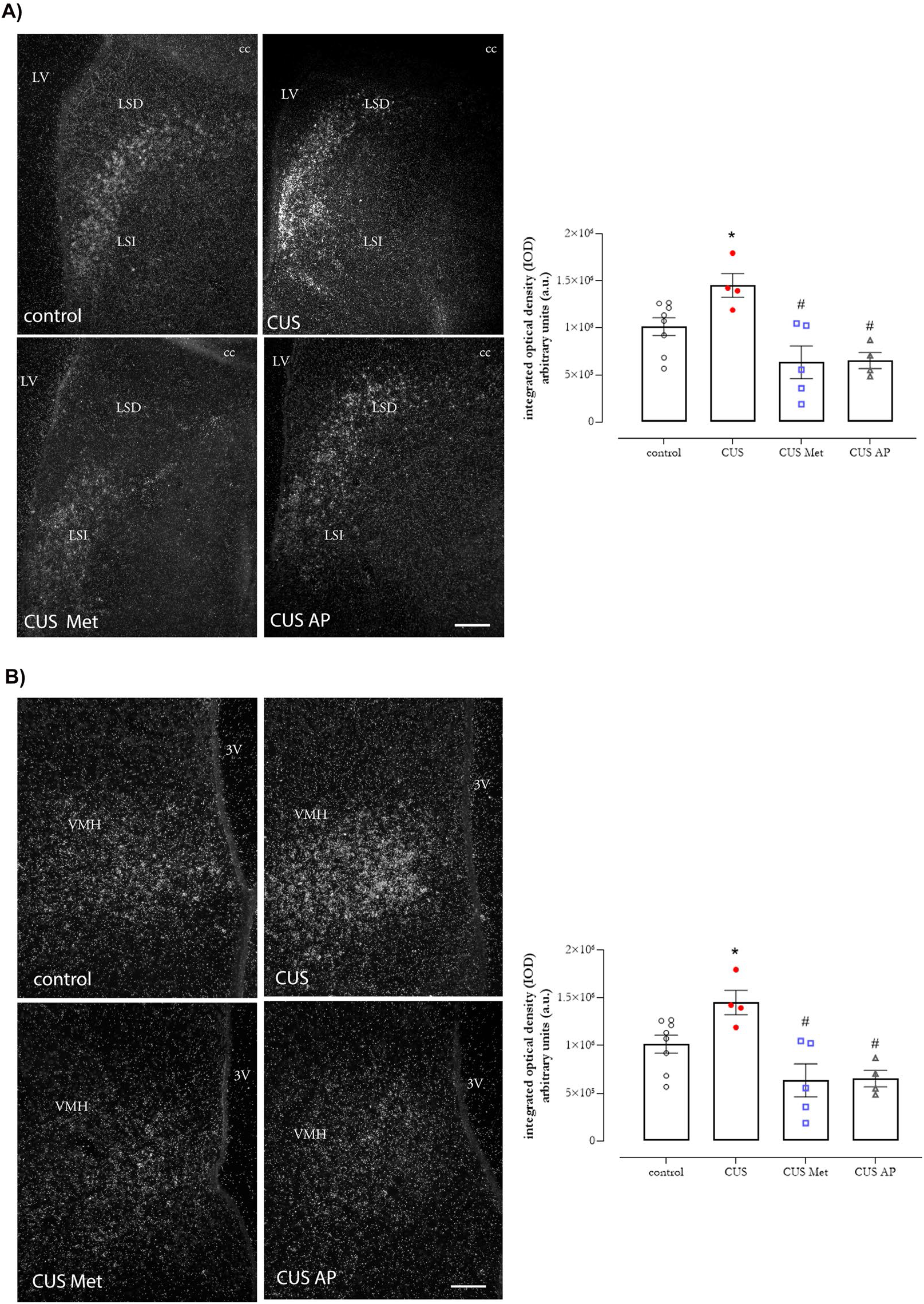
CUS-induced increase in the CRF_2_ RNAm expression in hypothalamic and extra-hypothalamic brain areas are modulated by GCs release and NE signaling. Representative photomicrography of CRF_2_ RNAm expression by *in situ* hybridization assay and the in the lateral septal nucleus intermedia part (A) and the ventromedial hypothalamic nucleus (B) in control and 14 days of chronic unpredictable stress (CUS) exposed animals treated or not with metyrapone (Met, 50 mg/kg, i.p., 30 min before each stress session) or atenolol and phentolamine (AP, 5 mg/kg and 2.5 mg/kg respectively, i.p., 20 min before each stress session) at medial levels of LSI and VMH (0.20 and −3.14 mm from Bregma for LSI and VMH, respectively). The graphics illustrate the quantification of the integrated optical density (IOD) represented in the panels. Results are presented as mean ± SEM. Significance differences between groups are indicated as * p < 0.05 vs control; # p < 0.05 vs CUS, Kruskal Wallis and Mann Whitney tests (n = 6 for each group). Abbreviations: LSI - lateral septal nucleus intermedia part; LSD - lateral septal nucleus dorsal part; LV - lateral ventricle; cc – corpus callosum; VMH – ventromedial hypothalamic nucleus; 3V – 3rd ventricle.

**Table 1:** CUS-induced increase in the CRF_2_ RNAm expression in hypothalamic and extra-hypothalamic brain areas are modulated by GCs release and NE signaling. CRF_2_ RNAm expression was measured by *in situ* hybridization assay the in the lateral septal nucleus intermedia part (LSI) and the ventromedial hypothalamic nucleus (VMH) of control and 14-days of chronic unpredictable stress (CUS) exposed animals treated or not with metyrapone (Met, 50 mg/kg, i.p., 30 min before each stress session) or atenolol and phentolamine (AP, 5 mg/kg and 2.5 mg/kg respectively, i.p., 20 min before each stress session). Results are presented as mean ± SEM of the integrated optical density (IOD, arbitrary units). Significance differences between groups are indicated as * p < 0.05 vs control; # p < 0.05 vs CUS, Kruskal Wallis and Mann Whitney tests (n = 6 for each group).

|  | CRF<sub>2</sub> mRNA expression |  |
| --- | --- | --- |
|  | LSI | VMH |
| Control | $1,0 \times 10^6 \pm 9,4 \times 10^4$ | $7,9 \times 10^5 \pm 1,0 \times 10^5$ |
| C Met | $6,6 \times 10^5 \pm 1,6 \times 10^5$ | $1,7 \times 10^6 \pm 4,2 \times 10^4 *$ |
| C AP | $6,6 \times 10^5 \pm 1,4 \times 10^5$ | $5,5 \times 10^5 \pm 1,2 \times 10^5$ |
| CUS | $1,4 \times 10^6 \pm 1,3 \times 10^5 *$ | $2,4 \times 10^6 \pm 2,2 \times 10^5 *$ |
| CUS Met | $6,4 \times 10^5 \pm 1,7 \times 10^5 \#$ | $6,4 \times 10^5 \pm 1,1 \times 10^5 \#$ |
| CUS AP | $6,5 \times 10^5 \pm 8,6 \times 10^4 \#$ | $1,3 \times 10^6 \pm 2,7 \times 10^5 \#$ |

Noradrenergic innervation is widespread in the CNS, and its targets include regions implicated in stress-induced effects. We also observed that NE signaling could modulate plasmatic CORT levels (Figure 1). Because of that, we tested whether the expected stress-induced NE increase contributed to the stress effects on CRF_2_ mRNA expression. We observed that the antagonism of both α- and β-adrenergic receptors (phentolamine and atenolol, respectively) immediately before each stress session reduced CRF_2_ mRNA expression in both LSI (Figure 2A) and VMH (Figure 2B) when compared to the CUS animals, 24 hours after the last stress session.

Chronic treatment with AP did not alter CRF_2_ mRNA expression in LSI and VMH in control animals, although the administration of Met (C+ Met, 1.7×10^6^ ± 4.2×10^4^ a.u) increased the expression of CRF_2_ mRNA in VMH when compared to control untreated animals (p < 0.05) (Table 1).

### 3.3. CUS persistently increased GR expression in brain regions related to GCs release control, an effect dependent on GCs and NE signaling

We evaluated the GR expression in several brain areas implicated in stress response. Our results showed that CUS increased the number of GR-immunoreactive cells in the LSI, PVH, CeA, and BST compared to the control group (p < 0.05). Unlike CRF_2_ mRNA expression, the influence of GC or NE in CUS-induced GR levels varied according to the nucleus analyzed. For the LSI and CeA, both treatments (Met or AP) prevented the CUS-induced increase in GR protein levels (p<0.05; Figures 3A and 4B). In the BST, only the AP treatment blunted the CUS-induced GR levels increase, suggesting the dependence of NE for this effect and not GCs (Figure 3B). In the PVH, the primary hypothalamic nucleus in controlling stress responses, both treatments did not prevent the CUS-induced increase in GR expression (p > 0.05) (Figure 4A).

**Figure 3.**
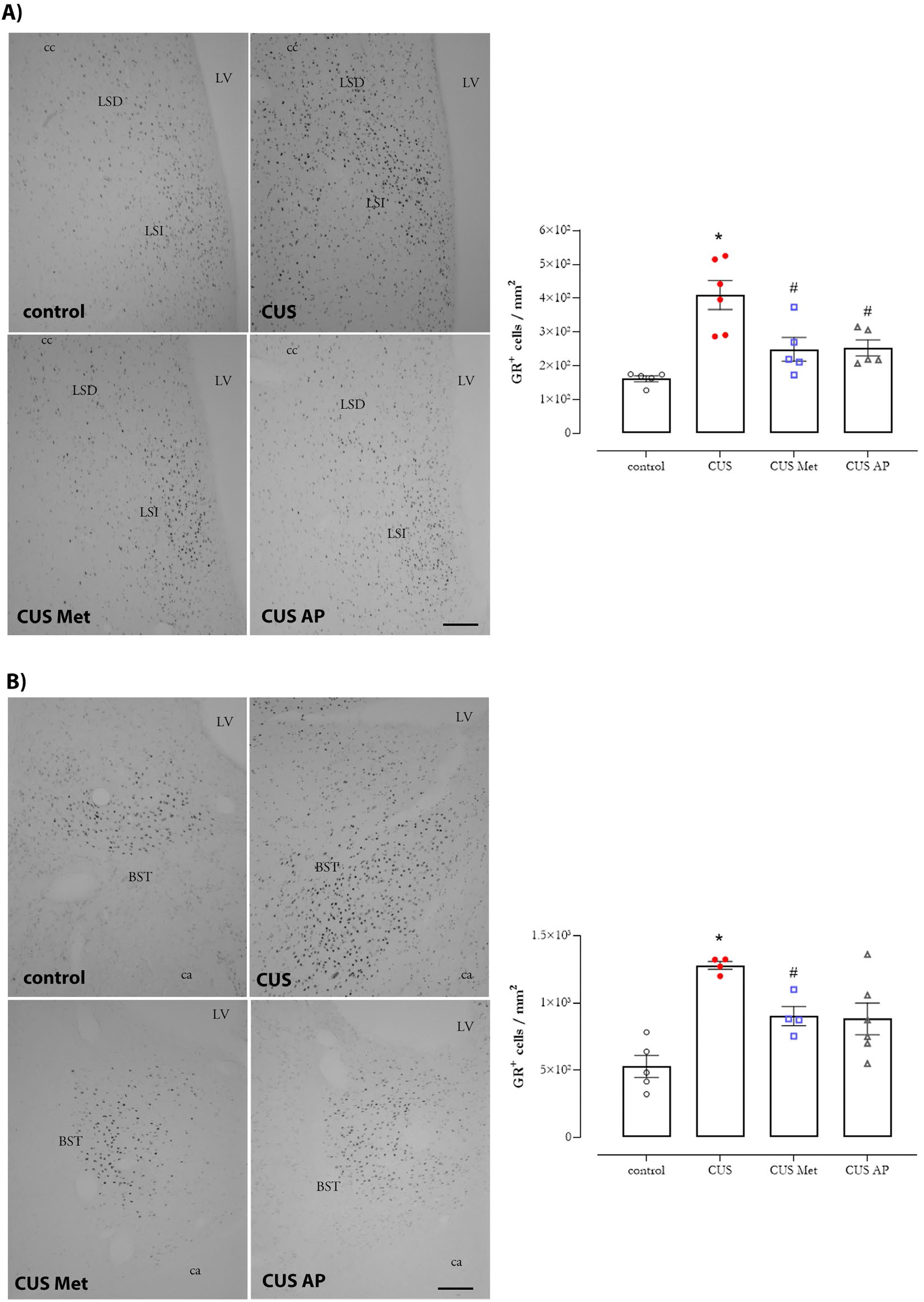
CUS-induced increase in the GR protein levels in hypothalamic and extra-hypothalamic brain areas are modulated by GCs release and NE signaling. Representative photomicrography of immunoreactive GR cells in the lateral septal nucleus intermedia part (A), bed nucleus of the stria terminalis (B) in control and 14 days of chronic unpredictable stress (CUS) exposed animals treated or not with metyrapone (Met, 50 mg/kg, i.p., 30 min before each stress session) or atenolol and phentolamine (AP, 5 mg/kg and 2.5 mg/kg respectively, i.p., 20 min before each stress session). The graphics illustrate the mean of GR immunoreactive cells /mm2. Significance differences between groups are indicated as * p < 0.05 *vs* control group; # p < 0.05 *vs* CUS group Kruskal Wallis and Mann Whitney tests (n=6 for each group). Abbreviations: LSI - lateral septal nucleus intermedia part; LSD - lateral septal nucleus dorsal part; LV - lateral ventricle; cc - corpus callosum; BST - bed nucleus of the stria terminalis; ca - anterior commissure.

**Figure 4.**
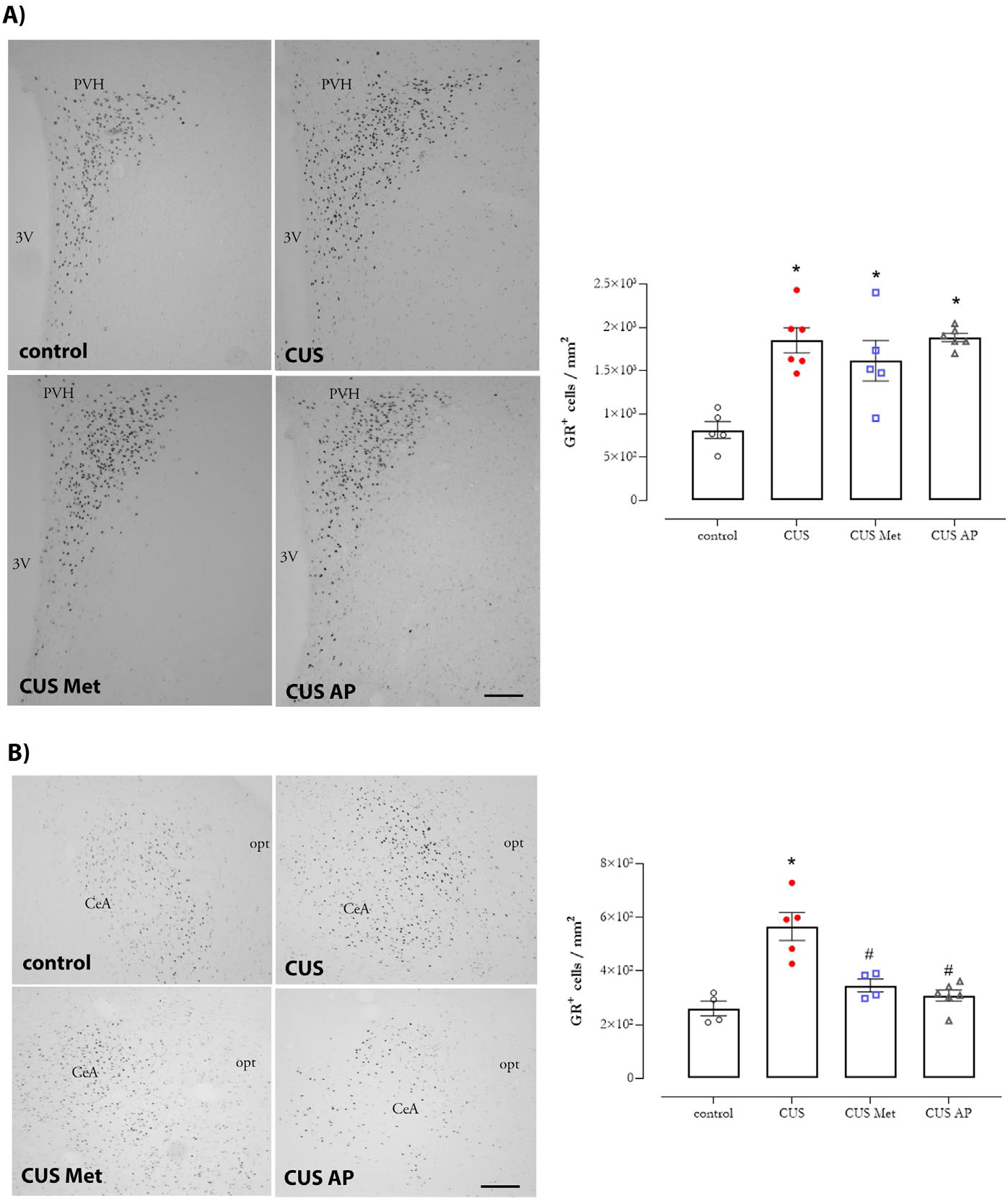
CUS-induced increase in the GR protein levels in hypothalamic and extra-hypothalamic brain areas are modulated by GCs release and NE signaling. Representative photomicrography of immunoreactive GR cells in the paraventricular hypothalamic nucleus (A), central amygdaloid nucleus (B) in control and 14 days of chronic unpredictable stress (CUS) exposed animals treated or not with metyrapone (Met, 50 mg/kg, i.p., 30 min before each stress session) or atenolol and phentolamine (AP, 5 mg/kg and 2.5 mg/kg respectively, i.p., 20 min before each stress session). The graphics illustrate the mean of GR immunoreactive cells /mm2. Significance differences between groups are indicated as * p < 0.05 *vs* control group; # p < 0.05 *vs* CUS group Kruskal Wallis and Mann Whitney tests (n=6 for each group). Abbreviations: CeA - central amygdaloid nucleus; opt - optic tract; PVH - paraventricular hypothalamic nucleus; 3V - 3rd ventricle.

Chronic treatment with AP did not alter the GR levels in control animals in any nuclei analyzed. However, MET treatment increased GR levels in PVH when compared to control animals not pharmacologically treated (p < 0.05) (Figure 3 and Table 2).

**Table 2.**
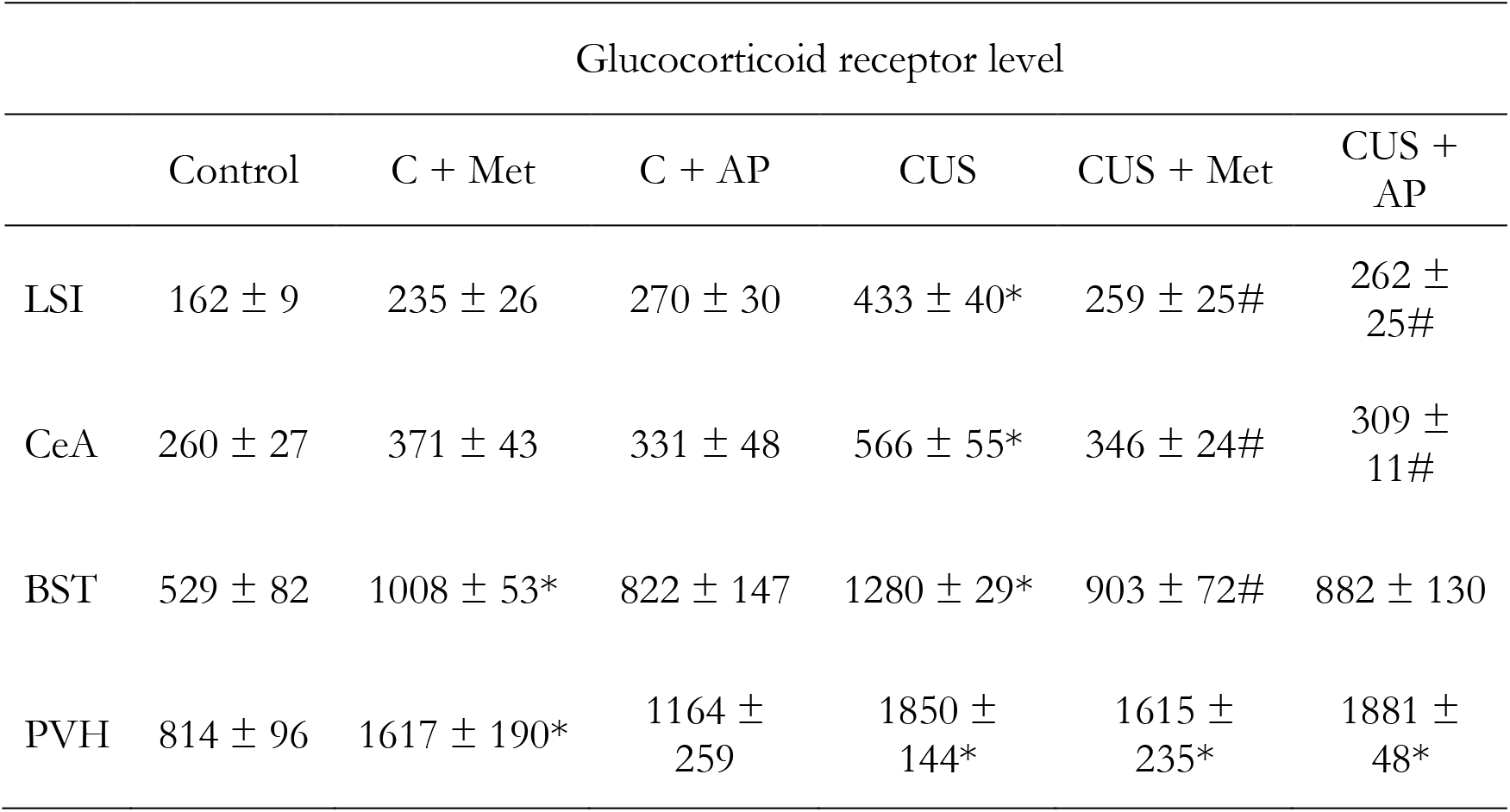
CUS-induced increase in the GR protein levels in hypothalamic and extra-hypothalamic brain areas are modulated by GCs release and NE signaling. The results illustrate the mean of GR immunoreactive cells /mm2 in the lateral septal nucleus intermedia part (LSI), central amygdaloid nucleus (CeA), the bed nucleus of the stria terminalis (BST), and the paraventricular hypothalamic nucleus (PVH) of control and 14-days of chronic unpredictable stress (CUS) exposed animals treated or not with metyrapone (Met, 50 mg/kg, i.p., 30 min before each stress session) or atenolol and phentolamine (AP, 5 mg/kg and 2.5 mg/kg respectively, i.p., 20 min before each stress session). Significance differences between groups are indicated as * p < 0.05 *vs* control group; # p < 0.05 *vs* CUS group Kruskal Wallis and Mann Whitney tests (n=6 for each group).

We next analyzed if the CRF_2_ mRNA expression and GR were present in the same cells in the rat brain. Our results showed both CRF_2_ mRNA and GR in the same cells of LSI (Figure 5).

**Figure 5.**
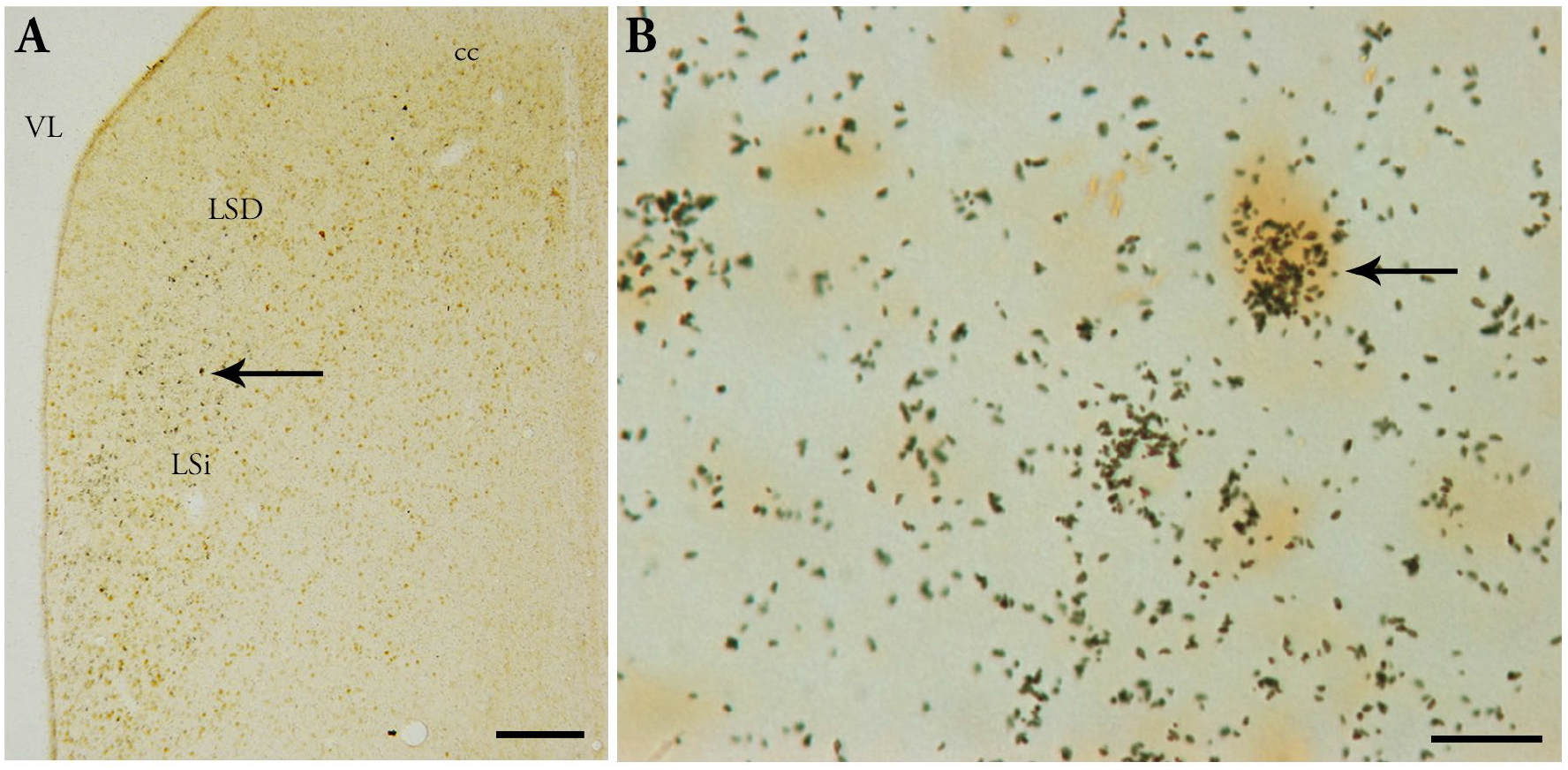
Representative photomicrography of co-localized CRF_2_ RNAm expression (silver grains) and GR (DAB, brown color) in the LSI cells indicated by the arrows. Abbreviations: LSI - lateral septal nucleus intermedia part; LSD - lateral septal nucleus dorsal part; LV - lateral ventricle; cc - corpus callosum. Scale bar: A: 100 μm; B: 10 μm.

### 3.4. CUS did not induce anxiety-like behavior when tested 24 h after the last stress session

Because all these CUS-induced molecular changes and the relevance of GR and CRF_2_ in modulating some behaviors in chronically stressed animals [33,34,23,17], we analyzed the anxiety-like behaviors in our CUS paradigm using elevated plus maze (EPM) and open field (OF) tests after 24h of the last stress session. In our CUS protocol, no anxiety-related behaviors were observed in CUS animals in both tests. In the EPM test (Figure 6A), no difference in the time spent in the closed (68 ± 7%, CUS) and open arms (16 ± 6%, CUS) were observed when compared to control animals (62 ± 7% and 21 ± 7%, respectively), evaluated 24 h after the last stressor. Consistently, in the OF, the CUS animals did not present anxiety-like behavior (Figure 6B) since no difference in the time spent in the central zone was observed in CUS group (95 ± 1.07 %) when compared to control animals (96.25 ± 0.72 %).

**Figure 6.**
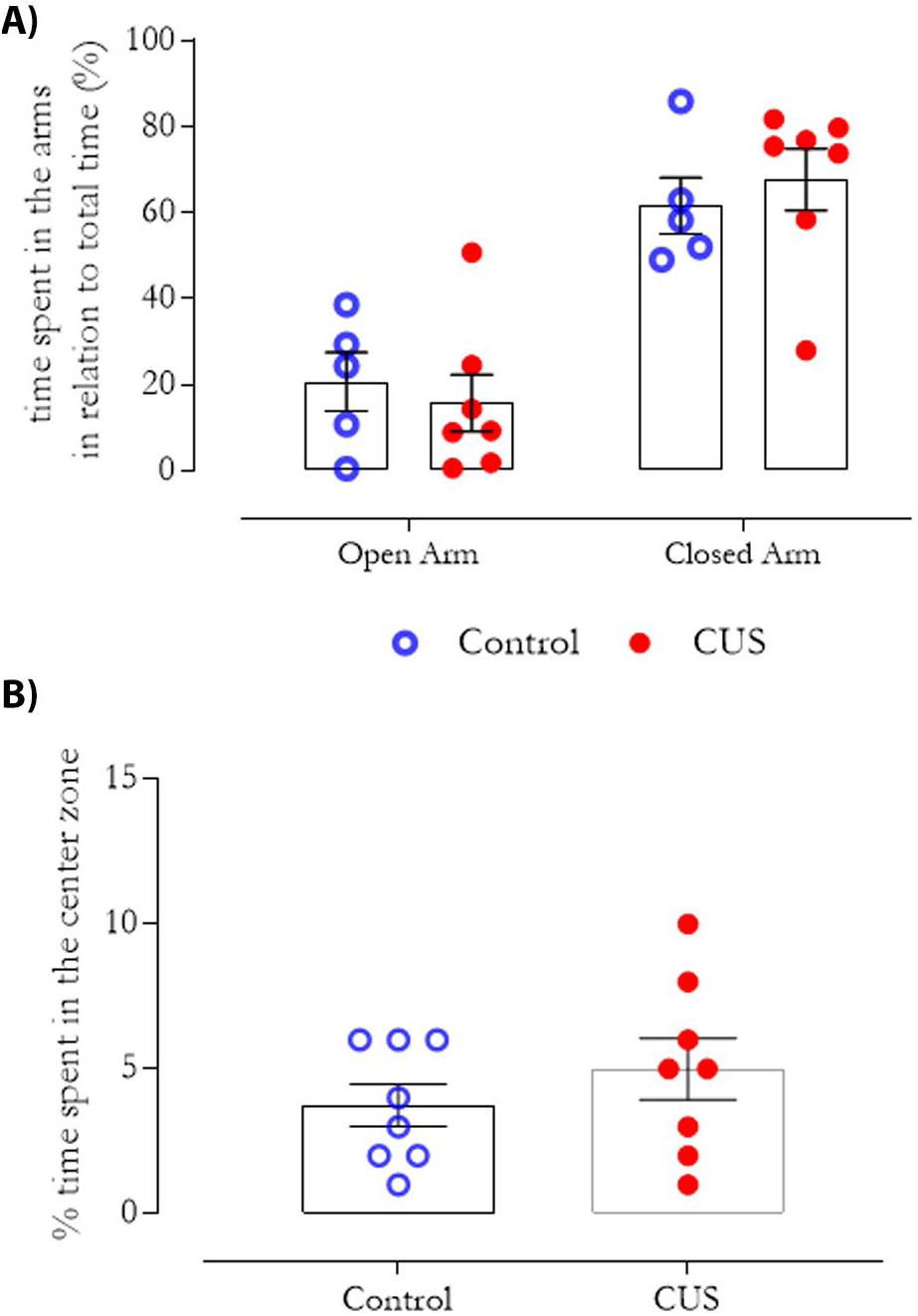
CUS did not induce anxiety-like behavior in rats measured by the elevated plus maze (EPM) and open field (OF) tests. Results are presented as mean ± SEM of the percentage of the time spent in open and closed arms in the EMP test (A) and the time spent in the center zone of the OF test (B) 24 hours after the last stress session. No significant differences between groups were observed. Student’s t-test, p>0.05; n=12 for EPM and n = 8 for OF.

## 4. Discussion

In the present study, we showed that 14 days of CUS increased plasmatic CORT levels that persisted for up to 24 hours after the last stress exposure. This long-lasting effect was also accompanied by the hypertrophy of adrenal glands, suggesting hyperactivity of the HPA axis in response to CUS. The HPA axis hyperactivity has been reported in different stress contexts, mostly in the chronic ones, and can be considered a risk factor for developing systemic and neuropsychiatry diseases, such as depression and anxiety [35].

CUS-induced increase of plasmatic CORT levels has been related to a vicious cycle, which maintains the stress brain circuitry activated via GC actions on extra-HPA brain nuclei, due, at least in part, to the loss of the GC-mediated HPA axis negative feedback loop [36]. In this situation, according to [37], other brain nuclei, such as the medial amygdala and pre-frontal cortex, become essential in the HPA axis activation, and the following could be happening: 1) changes in HPA axis signaling; 2) amygdala hyperactivation, and 3) prefrontal cortex hypoactivation. These hypotheses are not mutually excluding, and the control of synthesis and release of GCs could be a multifaceted system, not exclusively dependent on the activation/inhibition of the HPA axis [37]. In addition to GCs, the NE released by LC and NTS could act in brain nuclei such as the amygdala and PVH to maintain HPA activity [11,38–40,17]. To our knowledge, for the first time, these results showed that the rise and release of NE are as vital as GCs, during every single stressor, to the establishment of the CUS-induced HPA axis hyperactivation.

The importance of NE signaling in controlling HPA axis activation is well known for acute stress paradigms [41]. However, identifying its participation in maintaining chronic hyperactivation of the HPA axis is relatively recent [42,43]. Centrally released NE also plays an essential role in arousal and alertness [reviewed on [39]], and the dysregulation of the noradrenergic system could contribute to the susceptibility of stress-related illnesses. NE (systemically or centrally released) can excite CRF-containing cells in the PVH to activate the HPA axis. This effect is thought to be mainly mediated by α1-adrenergic receptors, although a role for β-adrenergic receptors has not been excluded. It is suspected that NE may activate CRF-containing neurons in extrahypothalamic regions, such as the CeA, LSI, and hippocampus, during stress. However, this warrants further investigations [10].

The interactions between CRF- and NE-containing neurons within the CNS are well established. Intracerebral administration of NE and adrenergic agonists can change the activity of CRF-containing neurons, and, conversely, administration of CRF can alter the activity of noradrenergic neurons [reviewed on [39]]. Here we evidence the relevance of NE in this paradigm since our results showed that atenolol and phentolamine modulate the plasmatic CORT, GR levels, and CRF_2_ mRNA expression in brain structures related to the HPA control in animals submitted to CUS. Considering that atenolol does not cross the blood-brain barrier [44] and phentolamine does [29], therefore, acting in the α-adrenergic receptors throughout the brain, we could imply the role of NE signaling peripherally and centrally in our study. Nonetheless, it is important to consider that stress can modify the blood-brain barrier permeability, allowing some atenolol to pass through it and to play a role centrally as well [44,45]. Hence, we can infer that stress-induced NE release could, indeed, modulate GC levels, contributing to hyperactivation of HPA axis and the CUS-induced increase in CRF_2_ expression and GR levels in the LSI. CRF acts through the CRF_1_ and CRF_2_ receptors, in the initial and recovering phases of the stress response, respectively [34,12,17]. The CUS-induced increase of CRF_2_ mRNA expression could be related to the recovery phase in the CUS animals, since 72 h after the last stress session, the plasmatic CORT levels and the adrenal gland hypertrophy were not observed (data not shown).

Glucocorticoids themselves provide the dominant-negative feedback loop onto the HPA axis acting through MR and GR, at least in part, in the circuitry originated in the forebrain [46]. In the present study, we observed CUS up-regulated GR levels in the LSI, PVH, BST, and CeA of the animals. All these nuclei are related to integrating the HPA axis and the limbic system [46]. Our results also suggest that the LSI could be one of the significant brain nuclei involved in this process since the CUS increased both CRF_2_ mRNA and GR levels, not only in the same nucleus but also in the same cell. The LSI also directly innervates the hippocampus, the BST, and VMH, which could result in the CUS-induced HPA hyperactivation observed 24 h after the last stress session, contributing to the hypothesis proposed by [37] regarding HPA axis activation in chronic stress.

The LC is one of the essential noradrenergic nuclei in the CNS directly activated by acute stress (reviewed in [47]). Our results suggest, for the first time, that in a chronic stressful situation, NE could act either systemically, by sympathetic activation, or centrally, by LS activation, in brain areas such as BTS, CeA and PVH (CRF-NE feedback), contributing to the maintenance of the stress vicious cycle [48,49]. Here, we showed that NE participates in the CUS-induced increase in GR levels in the LSI, reinforcing the LC involvement in the HPA axis modulation as proposed by [36].

Therefore, we hypothesized that the loss of HPA axis regulation observed in the present work could be, in part, due to the loss of inhibitory GABAergic projections from the BST to the hypothalamus, since the CUS-induced alterations in LSI and CeA, as well as in plasmatic CORT levels could block the BST-hypothalamus pathway. However, we cannot neglect the role of the CeA-PVH projection over the HPA axis activity [50,51]. This projection can be altered since we observed that CUS induced an increase in the GR levels in CeA, which could be via CORT and NE signaling, as our treatment with Met and AP revealed. The molecular mechanism behind this NE participation is not fully elucidated. Nonetheless, our results point to the importance of alpha and beta-adrenergic receptors, as well as the increase of GCs (Figure 7).

**Figure 7.**
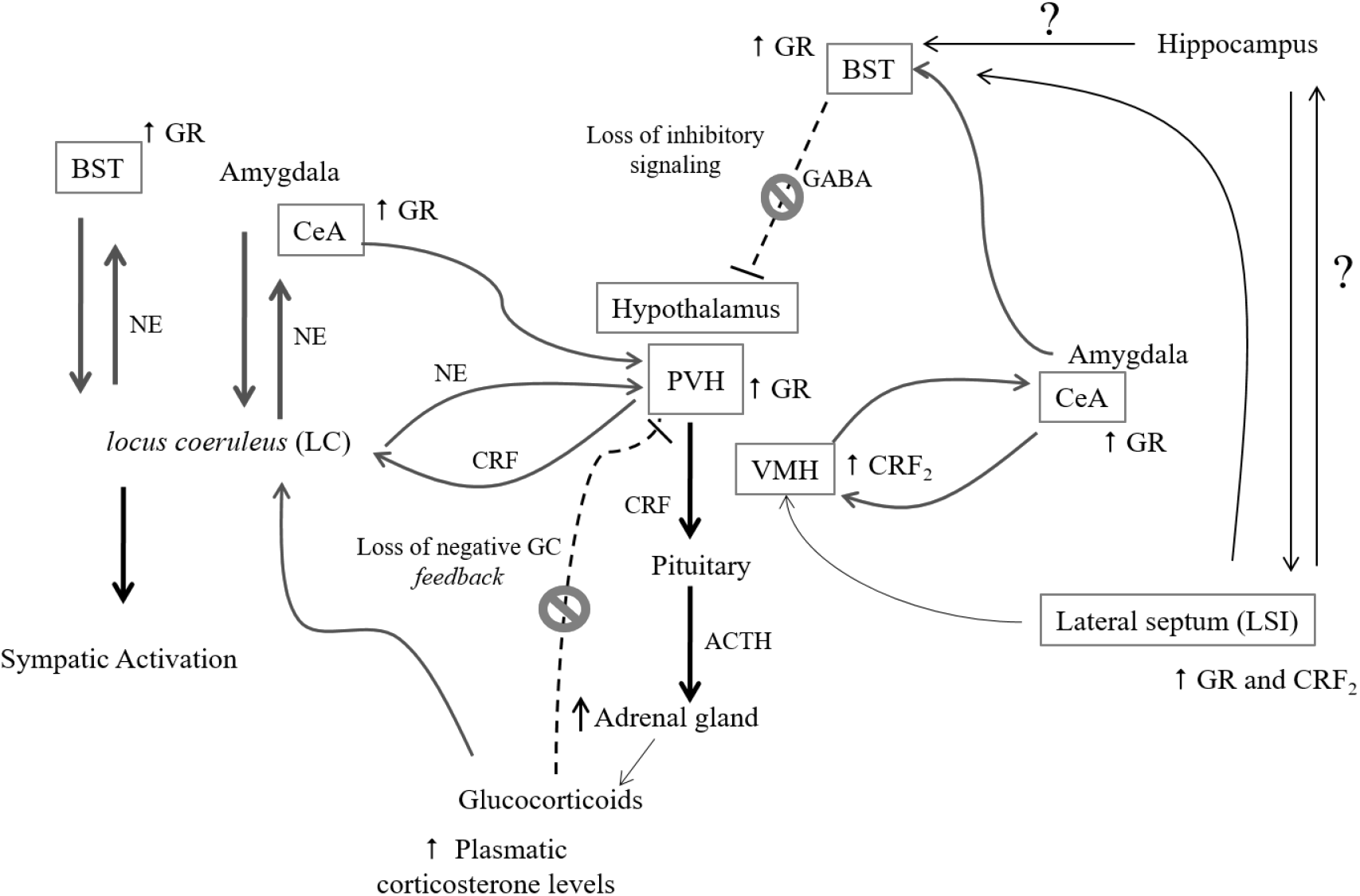
Schematic representation summarizing the main findings. Full lines represent excitatory signaling, while dotted lines represent inhibitory signaling. We hypothesized that CUS-induced increase in GR and CRF_2_ expression in LSI could be involved in the loss of HPA axis activity control, leading to sustained high CORT levels up to 48 hours after stress. These molecular CUS- induced changes modulate the excitatory LSI projection to VMH, BST, and hypothalamus. In addition, the loss of inhibitory signaling from BST to the hypothalamus could be the result of CUS-induced alterations in the BST projections from LSI, hypothalamus, and CeA.

Several studies have shown the participation of GR and CRF_2_ in specific brain regions such as the amygdala and lateral septal, in some behavioral alterations, including depression and anxiety-like behavior [33,34,23,12,17]. Despite the CUS-induced molecular plasticity in GR and CRF_2_, in the paradigm used here, this chronic stress protocol did not induce anxiety-like behavior in rat 24 hours after the last stress session. The anxiety-like behavior depends on CRF_1_ and CRF_2_ brain location, time, and intensity of stress [17]. Henry et al. (2006) tested the relation between CRF_2_ signaling and the LS with the anxiety-like behavior. Their data indicate that this effect depends on stress levels since a high-stress environment, but not lower stress conditions, induced anxiety-like behavior [52]. In the present work, CUS animals did not show anxiety-like behavior that could be associated with a combination of moderate stress conditions, recovery interval between each session, and social support since the animals were grouped in 3 to 4 animals per cage throughout the experiments. The effects of social support in grouped rats are attributed to neural plasticity and some neurotransmitters release such as oxytocin, well known for its anxiolytic properties [53,54].

Considering the absence of behavioral alterations and the increase of CRF_2_ mRNA in the present study, we also suggest a possible protective role of CRF and urocortins in this CUS paradigm. Some studies correlate CRF_2_ to the reestablishment of the stress response. They suggest that the CRF_2_ activation is responsible for ensuring physiological and psychological homeostasis and counteracts the initial stress-response-provoking effects and anxiety-like behavior induced by CRF_1_ activation [reviewed in [12]]. We also observed that this CUS-induced resilience-prone state lasted at least 10 days after the last stress session, including when a new challenge (2 h of restraint stress) was introduced (data not shown).

Prolonged or chronic stress changes the rules under which the body regulates homeostasis, requiring new strategies for successful adaptation. Only when the homeostatic pressure becomes too high, the body enters a state of genuine distress that can lead to morbidity and mortality. Thus, stress alters the physiological milieu over a long-term manner (adaptation through change, or allostasis), and the body’s response to these changes lies at the center of both successful stress resilience as well as its transition to pathology [55]. Therefore, while responses to chronic stress can clearly have negative consequences, one must consider the possibility that they may solve the organisms’ pressing problems at the expense of future success. The switch from adaptive to maladaptive stress responses will be heavily dependent on the individual’s constitution, based on genetic and acquired strategies to efficiency and limit overdrive of stress systems. In conclusion, the present study highlights that CRF_2_ and GR, mainly in LSI, play a role in fine-tuning CUS responses, which depend on GC and NE signaling. Although further studies are necessary to explore the mechanisms behind these results, we can suggest that CUS-modulated molecular changes improve animals’ ability to cope with stressful events, which results in a state of resilience for anxiety-like behavior.

## Declarations

## Acknowledgments

We gratefully thank Guiomar Wiesel and Larissa de Sá Lima for technical assistance.

## Availability of data and material (data transparency)

The data that support the findings of this study are available from the corresponding author, upon reasonable request.

## Code availability (software application or custom code)

Not applicable

## Authors’ contributions

M.B.M. designed research, performed research, analyzed data, wrote the paper, revised the paper final version; J.M. designed research, performed research, revised the paper final version; L.S.N. performed research, analyzed data, revised the paper final version; N.B.S. performed research, analyzed data, revised the paper final version; L.S. analyzed data, contributed unpublished reagents/analytic tools, revised the paper final version; R.C: contributed unpublished reagents/analytic tools, revised the paper final version; C.S. contributed unpublished reagents/analytic tools, analyzed data, revised the paper final version; J.B. contributed unpublished reagents/analytic tools, analyzed data, revised the paper final version; C.D.M.: designed research, analyzed data, contributed unpublished reagents/analytic tools, wrote the paper and revised the paper final version.

## Ethics approval

All animals were used following the standards of the Ethics Committee for Animal Use of the Institute of Biomedical Sciences/University of São Paulo (CEUA- ICB 102/06 and 75/05) and the guidelines of the Brazilian National Council for the Control of Animal Experimentation (CONCEA).

## Consent to participate

Not applicable

## Consent for publication

All authors have contributed significantly and agreed with the entire content of this manuscript.

